# Sequential Modelling for Medical Image Segmentation

**DOI:** 10.1101/2023.11.01.565075

**Authors:** Chun Hung How, Jagath C. Rajapakse

## Abstract

An image can be seen as a long sequence of tokens with spatial structure hence an image segmentation task can be treated as a sequence-to-sequence task. Existing attention-based segmentation models have incorporated an up-sampling module at the pixel decoder, such design assumed the backbone to have multiple feature scales. Furthermore, the existing models did not emphasize the ability of sequential modelling for multi-tasking. In this paper, we propose to model image segmentation task as a long sequence prediction task with an improvisation on MegaByte. As MegaByte efficiently reduced the self-attention cost using multi-scale structure known as global and local models, we train two MegaByte models as an encoder and decoder for multi-tasking. Our multi-task model allows different segmentation tasks to be trained simultaneously and inferenced with a task prompt. The encoder takes in the image sequence to encode them as a key while the decoder takes in the conditional input as a query. As decoder conditional input is usually sparser than the image, we use a relatively shallow decoder. We demonstrate our method on segmentation of intracranial haemorrhages from computed tomography (CT) head scans and show comparable performance to existing models. On a brain tissue segmentation task on CT images, our method outperformed the existing methods especially on tiny structure.

**Clinical relevance:** CT images are the first line of head scans when stroke or haemorrhage is suspected. Segmentation of intracranial haemorrhages (ICH) can provide clinicians with important measures of brain lesions to decide on treatment procedure or surgical decision. Brain and tissue volumes are indicative of neurodegenerative diseases. The proposed techniques can be used to segment ICH and brain tissues from CT head scans.

## I. Introduction

Fully convolutional neural networks have been effective in medical image segmentation tasks due to its spatial inductive bias. Recently, the use of transformer models in image segmentation tasks [1]–[3] has shown comparable performances in medical imaging tasks. Mask2Former [4] has been proposed as a universal backbone-agnostic architecture for segmentation tasks. Recently, SegViT [5] has demonstrated that with a simple non-hierarchical [6] vision transformer (ViT) [7] encoder, the encoded image features can be used to compute similarity between classes to achieve semantic segmentation. OneFormer [8] uses task tokens to unify multiple segmentation tasks. However, these designs are still unique and complex, and they do not work on multiple medical segmentation tasks concurrently.

In this work, we build a simple model that works with different types of segmentation tasks. Inspired by [9], we train a model that can perform segmentation task based on the given task prompt. We improvised on the recently proposed MegaByte [10] to produce conditional output on image tasks. While original MegaByte was proposed to have single encoder structure, we have improvised on a decoder network to learn general tasks. We performed data mixing strategy during multi-task training while treating image and target pairs from different tasks as independent samples in the same batch with respective task prompt. We showed that simply training pixel by pixel can achieve comparable performance to the best model especially on tiny structure with just single unified model.

## II. Related work

Transframer [11] trained a single autoregressive model to perform tasks like semantic segmentation, depth estimation, video synthesis. UViM [12] proposed to solve high dimensional problem in task like panoptic segmentation by tokenizing the image with VQVAE [13]. Mask2Former [4] built on top of end-to-end object detection model [14] has made different segmentation tasks like instance, semantic and panoptic segmentation to be easily trained with a Transformer based architecture. However, the unitary end-to-end model is usually proposed in brain imaging problem such as the ICH challenge [15]. Since there are other segmentation tasks in CT head scan, our proposed transformer-based model simplified the architecture to perform multiple segmentation tasks.

## III. Method

### A. Model Overview

The first module in a ViT model is the patch embedding layer. It is a mitigation to downscale long sequence into local features for cost efficiency in the attention and feed forward module. Therefore, there are also other approaches such as downscaling the image with a convolutional backbone. Recently, Yu et al. has proposed a modification to the transformer architecture called MegaByte [10] such that long text sequence is able to be fitted into the model without words tokenization. Their work applies on image domain differently from ViT which conventionally patching the image into smaller spatial resolution. Instead, there are two transformer modules in the architecture that designed for multiscale features. However, it is not directly applicable if we allow model to take in some conditions other than the input sequence. To train on multiple segmentation tasks on MegaByte, we adapted the architecture with a decoder following [16].

The architecture of our model is illustrated in Figure 1. The encoder learns the structure in the image domain and pass it to the decoder to return the correct format according to the task. More specifically, if we are solving for a task *y* that only conditions on the input image *x*, that is solving *p(y*|*x)* such as in semantic segmentation, the decoder input is set as an image-length sequence with values all zeros following the task prompt. On the other hand, if the task has an additional condition *c* on the image *p(y*|*x*,*c)* such as instance segmentation given a bounding box *c*, the condition is given as the decoder input but limited by the image sequence length. MegaByte exploited the advantage of using multi-scale feature to accommodate the long sequence into the model, which would be very costly with the classic transformer. In our work, we use only one global and one local level module in the model.

**Figure 1.**
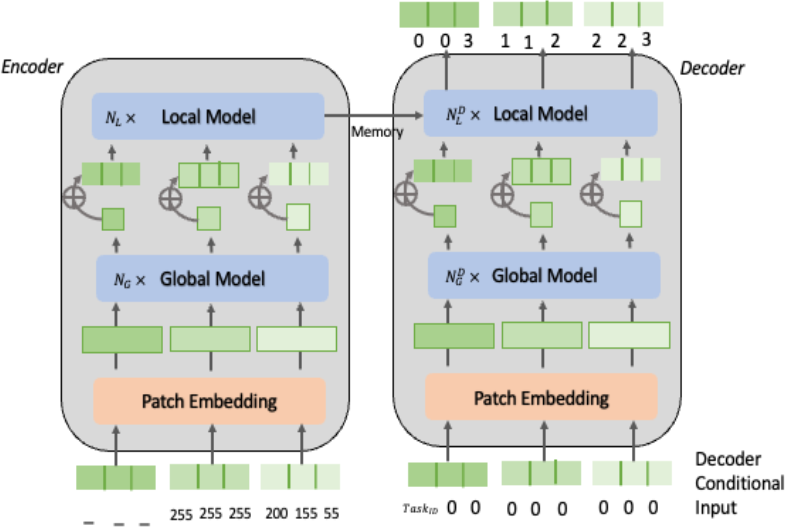
Our proposed architecture for segmentation tasks. The left module is the encoder like a transformer encoder module. The right module is the decoder module. Segmentation task is modeled as sequence-to-sequence task. The input to the decoder can be a single task ID token during inference.

First, in the encoder, we have a patch embedding module to project the tokens within a patch into a representation embedding for each patch. It works similarly like a convolutional down-sampling module, but these patch embeddings could attend to other patches globally. Next, each patch embeddings are appended as the first local token within the patch where the local model is responsible for attending tokens within the patch. Then, the final output representation of the encoder is used as the key (*memory*) and the embedded decoder input would be the query in the decoder local model. Since the decoder query needs to attend to all positions of encoder output, a causal mask was not used in the decoder but used only in the encoder. Note that the cross attention only occurs at local model which is patch-to-patch. This is because the global information has already been encoded into the patch. As for implementation, we followed the same model configuration as in [10] for the encoder. Since the information given in decoder input will be sparser as compared to the encoder, we train a relatively shallow decoder.

### B. Multi-task preprocessing

The input sequence to the decoder has a special token in the beginning that specifies the task. In our case, one for instance segmentation and another for semantic segmentation. Both of our segmentation tasks have 2D images as input, restricted to the same resolution universally. First, we flatten the image into a sequence by splitting the image into patches, and then perform raster scan within each patch as we encourage the local model to learn the attention of the neighbor pixels. We think that such spatial bias could help image segmentation as patch scan was shown to be superior to plain raster scan in image tasks. [10] In this way, the global model embeds each local patch and learn the attention across patches, which is not the case if we flatten them row by row and left to right. To generalize the decoder model to all kinds of inputs and conditions such as a 2D mask or bounding boxes, the decoder also takes in a sequence of the same size as encoder input but the rest of the unused elements in the sequence are filled with zeros.

For instance segmentation task, we give the quantized bounding box [9] on the region of interest. For semantic segmentation task, we simply give a zero vector with the size of the image. Both segmentation tasks return the pixel-classified masks as sequence as illustrated in Figure 2. Semantic segmentation task returns a sequence of integers representing the classes of each pixel while instance segmentation only has background and foreground classes.

**Figure 2.**
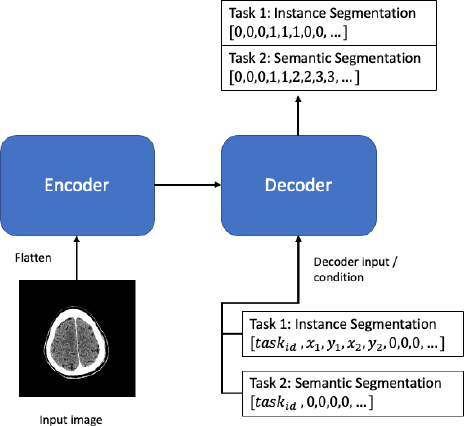
An illustration of the prediction sequence in training two segmentation tasks: instance and semantic segmentations. Following the task index, we gave quantized box coordinates for instance segmentation, or all zeros for semantic segmentation.

**Figure 3.**
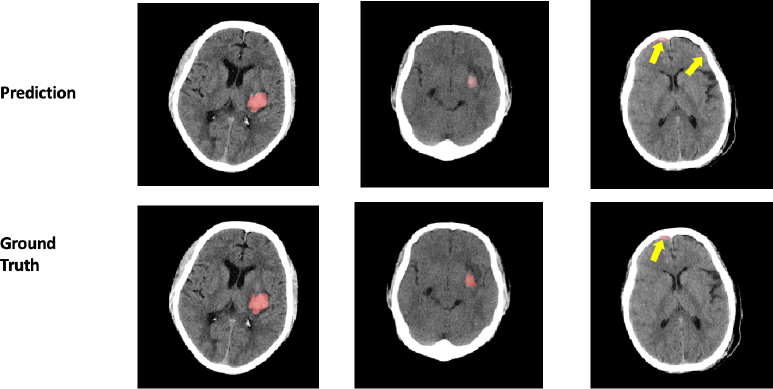
Examples of ICH segmentation from our model. The ground truth box that localizes the lesion is given to the decoder as input. Top row shows predictions, and the bottom row shows ground truths.

## IV. Experiments

In what follows, we discuss the datasets, setup, and results of our experiments. The performances of our model in comparison to existing methods are presented. Note that the datasets we used here are all head CT scans and the segmentations were specifically performed on 2D images.

### A. Datasets

#### A.1. ICH Lesion Segmentation

We obtained publicly available datasets from INSTANCE’22 challenge [15], [17] and PhysioNet [18]. The former has 893 slices from 100 scans that contain annotated intracranial hemorrhage (ICH) lesions. The latter has 273 slices from 36 scans that contain annotated ICH lesion. We obtained the bounding boxes that locate the lesions using PyTorch mask-to-boxes function on the ground truth masks.

#### A.2. Brain Tissue Segmentation

To expose the model to learn more about brain anatomical structure while training lesion segmentation, we introduced brain tissue segmentation task to the model. The model is trained to segment brain tissues: grey matter (GM), white matter (WM), Cerebrospinal fluid (CSF); and background from head CT scans. We used the head CT images of 37 subjects in CERMEP-IDB-MRXFDG [19]. The second dataset we used is from OASIS-3 [20] containing 305 subjects. We obtained their brain tissue ground truth mask using FSL [21] on the MRI images and overlayed on the registered CT. For OASIS-3 datasets, we performed brain extraction task.

### B. Results

In the experiment, we used 6 layers for global and local model in the encoder and 2 layers for the global and local model in the decoder. The number of parameters of encoder and decoder are 11.2 million and 8.5 million respectively. We trained both tasks in a data mixing fashion [9] with image size 256 for 50 epochs parallelly using 3 nodes of NVIDIA A100 80GB with batch size 12, sampled evenly across each task and node. We set 10 epochs warm up learning rate to 0.001, and applied StepLR to gradually reduce learning rate to 0 across the rest of the epochs. We applied mild random augmentations such as scale (0-5%), translation (0-10%), rotation (−15°,15°), shear (−5,5), gaussian noise on the training set.

#### B.1. ICH Lesion Segmentation

The model is trained on lesion segmentation given 4 coordinates of the bounding box. The decoder input sequence starts with a task index and the quantized coordinates [9], followed by all zeros. However, the bounding box information is given to ease the task complexity, and there is no restriction on pixel classification outside of the box. We compare to two other models, SwinUNet [3] and TransDeepLab [2], which were trained solely on the same training set. These two models were trained for 300 epochs with random augmentations to detect and segment the lesion in the images. We used INSTANCE’22 for the experiment, we randomly chose 10 of the subjects (10%) and assigned all the annotated slices (73) in their scans as test set. The average dice scores of segmentations are given in Table I. As seen, we achieved decent performance comparing to existing model.

**TABLE I.**
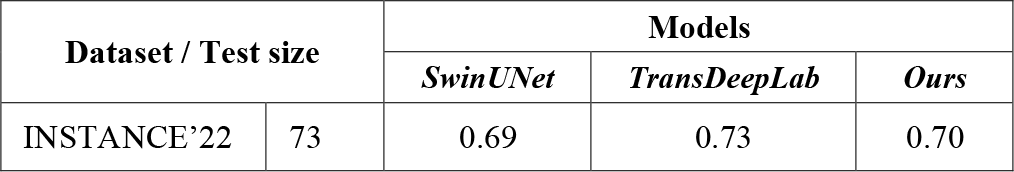
ICH leson segmentation test dice scores.

#### B.1.2. Generalizability

We performed a test using the trained model on unseen PhysioNet dataset. We used the entire PhysioNet dataset as a general test set since it is a relatively small dataset. Our hypothesis is that the images from PhysioNet has similar quality and scanner setting as INSTANCE, therefore our trained model should be able to perform similarly. For this analysis, we categorized PhysioNet dataset by their lesion size (small, medium, large) according to 25^th^ percentile and 75^th^ percentile of INSTANCE lesion distribution, where small is below 25^th^ percentile, medium is between 25^th^ and 75^th^, and large is above 75^th^ percentile. We observed that PhysioNet dataset has higher proportion of small and medium lesions, leading to the degradation of generalization performance. Our result showed that the generalizability performance of our model dropped 40% overall from the test performance of training domain. We also observed that our model did not perform as good in large lesions but handled decently for small and medium lesions as shown in Table II.

**TABLE II.**
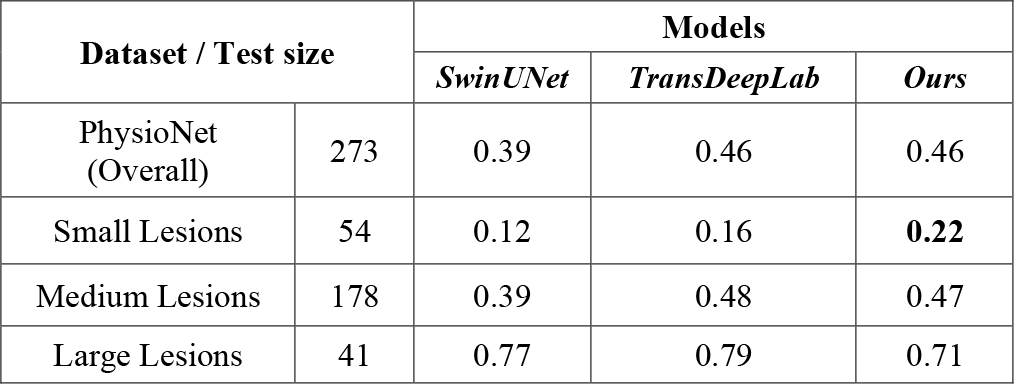
Generalisation test of leson segmentation on physionet (dice scores)

### B.2. Brain Tissue Segmentation

We trained the model to segment grey matter (GM), white matter (WM) and Cerebrospinal fluid (CSF). In total, there are 4 classes including background label. We picked 10 subjects from CERMEP-IDB-MRXFDG as test set. As this is a discriminative task with deterministic output, there is no sampling step during train nor test. We feed a sequence with the task index followed by all zeros therefore it learns to map the pixel class from the encoder output. For comparison, we used the classic UNet [22] and TransUNet [1] to train on the same train set. For this task, we additionally trained and tested on OASIS-3 solely using our model. Our model outperformed the other two models on tissue segmentation task as shown in Table III. We showed the examples of both datasets in Figure 4 and 5.

**TABLE III.** Test dice scores on brain tissue segmentation.

| Dataset / Test size |  | Class | Models |  |  |
| --- | --- | --- | --- | --- | --- |
|  |  |  | UNet | TransUNet | Ours |
| CERMEP-IDB-MRXFDG | 1720 | GM | 0.49 | 0.65 | **0.66** |
|  |  | WM | 0.44 | 0.66 | **0.67** |
|  |  | CSF | 0.20 | 0.53 | **0.55** |
| OASIS-3 | 8310 | GM | 0.51 | 0.62 | **0.63** |
|  |  | WM | 0.32 | 0.61 | **0.63** |
|  |  | CSF | 0.30 | 0.54 | **0.56** |

**Figure 4.**
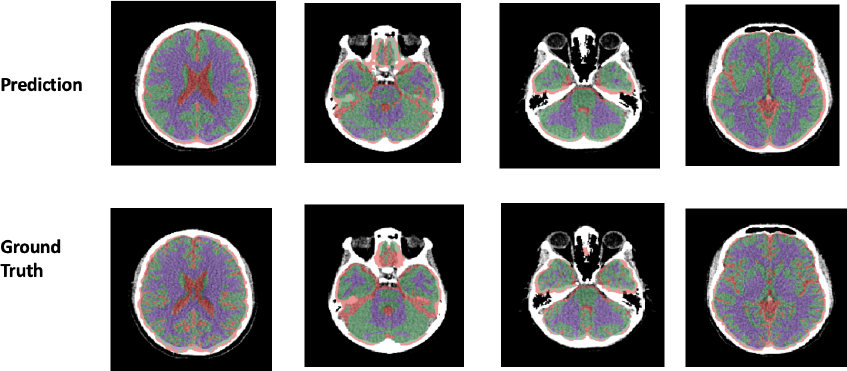
Test examples from CERMEP-IDB-MRXFDG in brain tissue segmentation task with our model. (GM – green, WM – purple, CSF – red)

**Figure 5.**
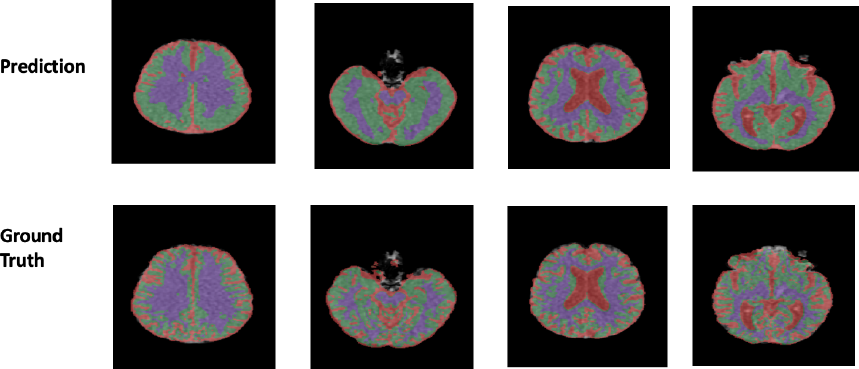
Test examples from OASIS-3 in brain tissue segmentation task with our model. (GM – green, WM – purple, CSF – red)

## V. Conclusion

Our unified multi-task model can be trained to answer different segmentation tasks with a task prompt. Our model achieved comparable or better results in ICH and our model outperformed the other unitary models on CT brain tissue segmentation which is heterogenous. Overall, our model is better in segmenting tiny and granular region.

## Notes

### Competing Interest Statement

The authors have declared no competing interest.

